# Foraging specialization and body size in seabirds

**DOI:** 10.1101/2023.06.25.546462

**Authors:** Juan Hernandez, José Ignacio Arroyo

**Affiliations:** Department of Ecology, Pontifical Catholic University of Chile, Santiago, Chile; Santa Fe Institute, Santa Fe, New Mexico, USA; Center for Mathematical Modeling, University of Chile and IRL-CNRS 2807 Santiago, Chile

**Author notes:** These authors contributed equally to this study.

## Abstract

Body size affects many biological processes since it predicts traits, timing, and biological rates. Some of these relationships are explained by the metabolic theory of ecology, which predicts that they should scale according to a power law with exponents multiples of 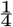. Here we study the relationships between foraging specialization, particularly the number of dietary categories and prey capture strategies, and seabird species size, based on a database of 342 species (representing more than 95 % of all species). In our analysis, we found a negative relationship between the number of dietary categories and the number of capture strategies with body size with exponents of -0.83±0.31 and -0.76±0.06. To explain these relationships in terms of first principles, we developed a simple model to explain the origin of this scaling based on well-established ecological scaling relationships. Our study suggests that foraging specialization is constrained by the energy used by an organism, providing a basis for future theoretical developments.

## Introduction

The body size of organisms affects their biological rates, times and many other traits, including physiological, ecological, behavioral, and evolutionary [1–7]. The relation of organism’s traits with their body size can be described by a simple mathematical relationship known as a scaling relationship [1–13],

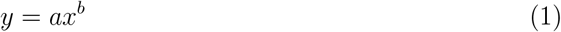

Where Y is a biological quantity, X is the organism’s body size whereas, *a* and *b* are constants. Commonly Eq. (1) can be expressed in log-scale, *log*(*Y*) = *log*(*a*)+*blog*(*X*), resulting in a simple linear relationship where ln() is the intercept and is the slope. The exponent has a simple interpretation. If the values are less than 1 (sublinear) it means that when the size increases the double, the Y variable increases less than the double. If the exponent is linear, is size increases the double the Y variables increase the double, and if the slope is greater titsn one (superlinear), the Y variable increases more than the double. Examples of sublinear scaling include metabolic rate [2], linear the size of different organs [14], and super linear the glucose uptake of different cancers [10]. It Is worth mentioning that exponents greater than 1 are common in systems such as cities, where the superlinear phenomenon indicates that the larger the better or worse, e.g. if the size of a city doubles, the per capita income or crime increases more than the double [15]. These relationships are commonly called ”economies of scale”, in the sense that as the size increases, costs decrease, and benefits increase.

Different models and theories have been suggested to explain the origin of these relationships, among which the metabolic theory of ecology (MTE) stands out. The MTE is a broad theoretical framework that initially emerged as a model explaining the relationship between body size and various organismic rates, including metabolic rate [3]. The original model argues that organisms transport matter through networks, such as the cardiovascular system, and based on some physical principles of the structure and dynamics of these networks a mathematical explanation emerges for the scaling of many traits and rates, all with exponents multiples of 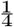, due to the fractal properties of biological architecture [16]. Later, principles of chemical kinetics would also be included to include temperature effects as well [2]. Subsequently, the theory was extended, and theoretical predictions were derived for several other variables at different levels of organization, from the individual to the ecosystem, such as molecular evolutionary rate, generation time, population growth rate, and species richness, among others [17]. Many of these predictions come from the integration of different principles for which there is good theoretical and/or practical support.

Despite the numerous predictions of the metabolic theory, there are still other possible developments, as metabolism (dictated by size) affects all biological processes. Here we focus on ecological specialization (also called niche breadth; [18], particularly forage specialization, which can be defined as what a species eats and how it eats it. The importance of understanding the causes and consequences of niche breadth, or degree of ecological specialization, is that it can affect other ecological traits such as the ability to colonize new environments (expand its geographic range), extinction risk [19], and ultimately to diversify [20–22]. The hypothesis that attempts to explain the relationship between dietary niche breadth and body suggests that size (which we will call “mechanical constraint” hypothesis) species that are small are constrained to handle only small resource items, whereas larger ones can handle small to large resource items, then they can access not only a bigger diversity of sizes but bigger diversity of prey species. Accordingly, it is expected that the diet breadth be bigger in larger body-sized species [23, 24]. Also, body size scales with home range [25] and it would be expected that species with a larger home range would have access to a greater amount of resources. There is no hypothesis suggested for the number of behaviors and body size.

Here we assess these relationships in seabirds, taking advantage of the availability and good quality of data for this group. Seabirds are a functional group and refer to all birds that live in the ocean environment, including marine and coastal. Seabirds are a paraphyletic group (including the orders Procellariiformes, Charadriiformes, Sphenisciformes, Pelecaniformes, Phaethontiformes, and Suliformes) and have independently and convergently acquired different adaptations to pelagic and aquatic life [26]. The study of their ecology [27] and particularly diet and foraging [28] is a relevant topic of study due to their relationship with humans, role in fisheries, and as indicators in marine environments. Our results show an inverse relationship between the foraging niche breadth with body size. As part of our discussion, we suggest a mechanistic hypothesis and a minimal (zero-order approach) model that could explain the emergence of these patterns.

## Methods

### Data

We quantified what a species eats and how it eats it by compiling data on the number of diet categories (e.g. fish, crustacean, cephalopods, etc.) and the number of prey capture behaviors a species have (scavenging, pursuit diving, or bottom feeding, surface seizing, etc.). These data were manually obtained from the Handbook of the Birds of the World Alive (http://www.hbw.com/). We used as an initial reference for diet guilds the classification of foraging guilds of North American breeding birds [29, 30]. We found a total of 29 strategies: scooping, wading; filtering or probing Foraging, ground forager, gathering, surface seizing, surface picking, deep plunging, shallow plunging, pursuit plunging, pursuit diving o bottom feeding, scavenging, pattering, hydroplaning, kleptoparasitism, contact dipping, hover dipping, dipping, aerial pursuit, shallow dives, wing, surface plung, kick splashing, battering or drowning, plunge diving, skimming, walking, surface follow, cannibalism, ship follower. Diets were categorized in 20 categories: carrion, reptiles, fish, cephalopods, crustaceans, echinoderms, insects, lampreys, tunicates, porifera, mammals, algae, worms, eggs from other species, plants, hydrozoa, chetonata, amphibians, plankton, polychaetes. It is worth mentioning that the data on the number of diet categories and prey capture behavior categories correspond to the known total accumulated number across the whole life cycle of a species, which includes any diet or strategies consumed or used during early and late ontogenetic development, and reproductive and non-reproductive period. Data on body size were collected from *BirdLifeathttp* : *//datazone*.*birdlife*.*org/home*. The dataset included 342 species. The number of diet categories present in species varied from 1-11 and strategies from 1-9. Body size ranged from 15.5-32500 g.

### Statistical analysis

Different approaches can be used to analyze the data; i) analyze the data for each single species, ii) average size by diet and category, or iii) categorize sizes into intervals and calculate the average of diets and categories for those categories. We took the second option as it makes sense in terms of prediction, but also report the other analysis in the supplementary material. We estimate the average body size for each nominal value of the number of strategies and diets. We fit the data to a power law such as the one in equation (1), but in logarithmic scale; *log*(*Y*) = *log*(*a*) + *bln*(*X*), using simple linear regression (model 1) by the least squares method, as implemented in the “lm” function of the R language (R Core Team).

## Results

We found a negative relationship between the number of diets a species has and their average body size with an exponent of -0.83±0.31 (intercept=3.33, r-sq.=0.44, p=0.025). The relationship between the number of strategies and their average size had an exponent of -0.76±0.06 (intercept=2.89, r-sq.=0.97, p=1*10-5). There was a positive relationship between the number of strategies and diet, with an exponent of 0.51±0.13 (intercept=0.14, r-sq.=0.85, *p* = 0.0000408). We also studied the relationship between the ratio number of strategies over the number of diets and the average body size, which had an exponent of -0.52±0.13 (intercept=0.14, r-sq.=0.23, p=0.0002). The relationship of number strategies over the number of diets is analogous to the cost/benefit ratio in finance, i.e., how many behaviors were used to get a given number of resource items. The general interpretation of these parameters is straightforward; for each gram of weight, a species does not eat 0.83 new diets or does not use 0.76 new strategies. It could also be interesting to think of this relationship inversely; for each new strategy a species losses (1/0.76=) 1.31 grams in size and for each new diet a species losses (1/0.83=) 1.2 grams in size, respectively. We also analyzed the data alternatively (see Methods), and obtained similar results; i.e. negative relationships between foraging specialization variables and size (Supplementary Material).

## Discussion

Previous studies have reported negative [31, 32], positive [23] or constant (i.e. invariant) [31] relationships between body size and diet niche breadth in herbivorous insects, terrestrial birds, and marine predators. Commonly, these papers relate each species to its diet niche breadth. Here we took a different approach to previous studies in that we use body size averages for different numbers of categories of strategies or diets used by species. These empirical relationships can help predict the number of different diet categories or prey capture strategies a given species can have.

Three points deserve discussion. First, one of the reasons why the mechanical constraint hypothesis might not be general is because many species capture their prey in groups, so individual size is not a constraint [33] as big prey could be captured by a group of small preys.

Second. It is worth mentioning that the patterns we found could change depending on other conditions such as the environment [34], e.g., in more productive areas such as near the tropics, could be a higher diversity of species, and consequently birds could have a higher diversity of diet categories available [35–37]. Also, it is important to have in mind that the number of diets and strategies change along the ontogenetic development so there could be an intra-specific scaling of size and number of diets as an organism [37]. Here we just focused on the accumulated number of diets and strategies across the lifespan, so other sampling approaches could take to slightly different parameter values for these relationships.

Third, which existing model explains the origin of these relationships? Different generalizations explain allometry, such as the model of exponential evolution of traits that gives rise to scaling laws [38, 39]. Briefly, this model suggests that traits could develop and/or evolve exponentially, and two exponential behaviors give rise to a power-law, which is observed in many traits with body size. In this model, the exponent would correspond to the ratio of the evolutionary rates of both traits. Below we outline a minimal model to explain the origin of the negative relationship between diets, strategies, and body size. This model is not a detailed explanation in terms of how multiple processes that are determined by size ultimately affect the foraging specialization but contains the minimal constraints that could explain it. This model is based on the hypothesis based on metabolic theory, which would predict a negative relationship, based on an evolutionary and physiological mechanism. The extent of resource use and the strategies to obtain them is a complex trait. Its evolutionary origin involves changes from the genomic level [40] to behavior. In general, evolutionary (and diversification) rates are faster in smaller organisms [41], so smaller animals would be expected to have evolved a greater number of diets and strategies.

### A minimal general model

The general derivation for the relationships between the number of diets and strategies and body size is that size affects rates of evolution molecular evolution, and if we assume that the rates of phenotypic evolution that ultimately determine the evolution of complex physiology and behavior that ultimately allows a species to be able to process a broad range of resources and develop different behaviors to obtain them. So, we have that mutation rate (*u*) depends on body size [41] according to; *u ∝ M* ^*−a*^, where *M* is size and *a* is a parameter. We will assume that the rate of phenotypic evolution *p*, depends on the rate of molecular evolution [42, 43]; *p ∝ u*. Then replacing in the previous expression, we have

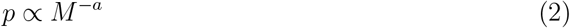

The phenotype includes not only all the physiological properties of organisms but also their behavioral properties, so with this simple model, we are able to explain the negative relationships between diet and strategies, and size. For example, in animals brain size, which is a proxy for different measures of intelligence, is a phenotype that ultimately affects behavior. Then it would be natural to think that phenotypes related to the brain and the nervous system, in general, are related to behavioral traits that ultimately affect even other traits. Just to illustrate how the mass dependence of the rate of phenotypic evolution ultimately can result in a mas-dependence of the transient or steady-state value of a trait, let’s assume a zero-order approach and say that the evolution of a trait X and size itself over a certain time interval changes as 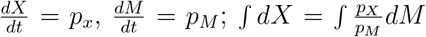. But 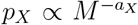 and 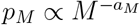, then

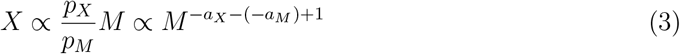

This simple model based on first principles explains the negative relationships of foraging specialization traits and body size and would allow us to predict the exponent values from simple scaling relationships. Some of the implications of the size-dependence of ecological specialization and foraging, in particular, is that it can link other traits that depend on foraging niche breadth such as range size [25], or extinction risk, diversification [44]. In birds for example, it has been showed that innovativeness is related negatively to extinction risk [19]. Given this simple theoretical basis, it is important to remark that despite here we have focused on a reduced functional group (seabirds), given the arguments exposed it would be expected to find that this is a general pattern in ecological systems. For example, if we think of bigger animals such as elephants or the blue whale, they are characterized by using a single strategy (grazing and filtering, respectively) and having a single diet (grass and krill, respectively) [45]. So, in future attempts, we will focus on expanding this database and testing the generality of this pattern in vertebrates or all animals and refining the model.

## Conclusion

In summary, we have shown a negative relationship between the number of diets, the number of foraging strategies, and the body size of seabird species and an invariant relationship between diet/strategy ratio (efficiency) and body size, with slopes of -0.83, -0.76 and -0.52. We have derived a simple general model to explain these patterns based on the size-dependence of molecular and morphological evolution and the evolution of traits as a linear dynamical system. This model is a generic system of first-order linear differential equations that models the dynamics of a couple of traits and accounts for the mass-dependence of evolutionary rates, based on principles of population genetics and empirical data. Our analysis and working model provide a basis for further theoretical refinements or extensions to explain the scaling to other ecological traits. For example, diet breadth is thought to affect other traits such as range, extinction risk, and diversification, which ultimately could be linked to size. Also, we hypothesize that this is a general pattern in animals, which will be a focus of future research.

**Figure 1.**
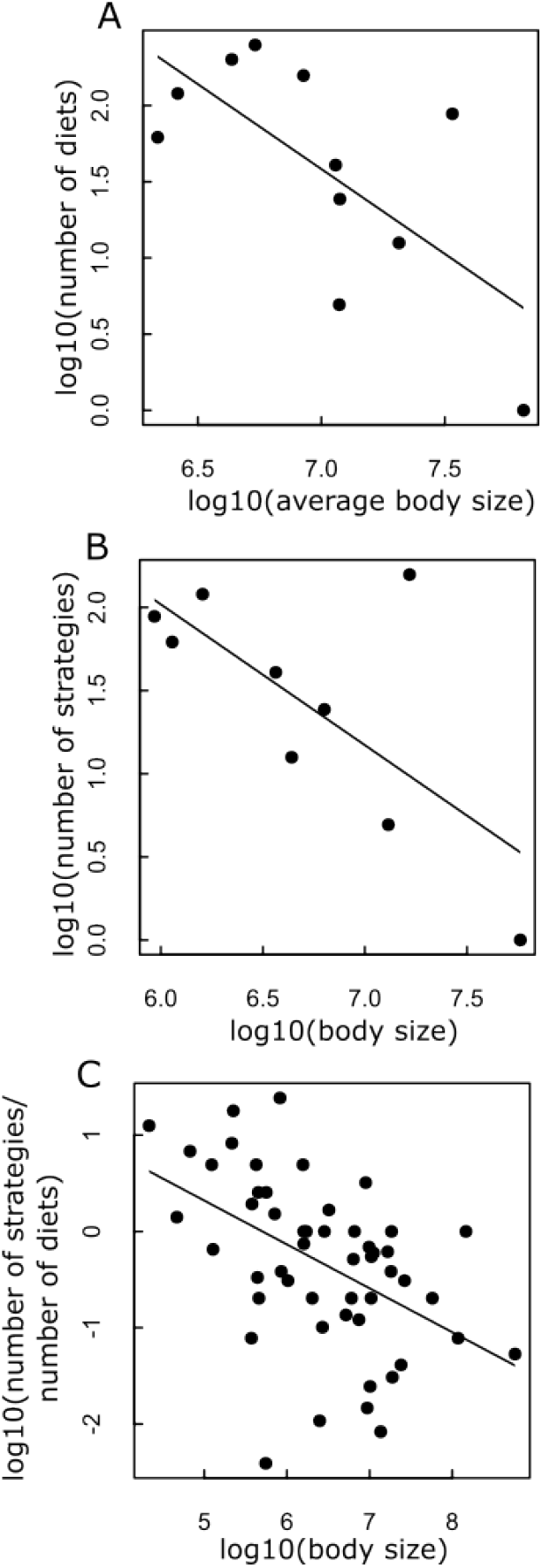
Relationships between (A) number of diet categories and average body size. Each dot in the plot is a set of species that consume the same given number of diet categories and their average body size. For example, the dot at the bottom left corresponds to a set of 19 species that eat one diet category (any of the listed in the Methods; Data section, not necessarily the same) and have an average of 2487.61 g of weight. (B) the number of prey capture behaviors and average-body size. Each dot in the plot is the set of species that have the same given number of strategies and their average body size. (C) the number of diet categories and number of prey capture behaviors and average body size. Each dot corresponds to the number of diets a given set of species have over their average number of strategies and their average body size. The slopes were −0.83, −0.76, and −0.52, respectively. See r-squared, and p-values in Table 1/Results section. Body size is in grams, axes are in log 10.

**Figure 2.**
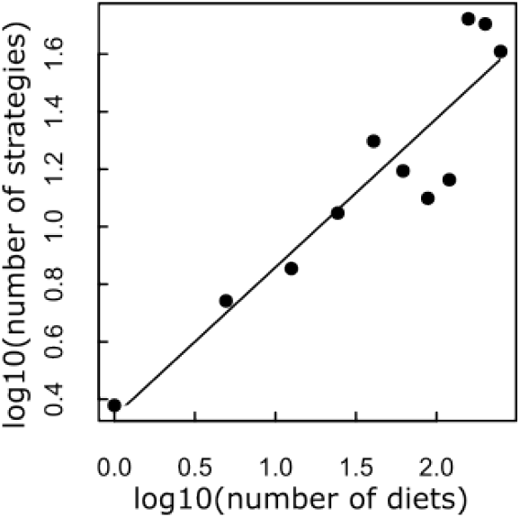
Relationship between the number of different prey capture behaviors and average diet categories a given set of species has. Each dot corresponds to the number of behaviors and the average number of diets for that number of behaviors. The slope is 0.51.

## Acknowledgements

JIA acknowledges a Beca de Doctorado Nacional (21130515) from the Chilean Agencia Nacional de Investigacion y Desarrollo (ANID) National Doctoral Scholarship, and USA NSF projects: ‘Building and Modeling Synthetic Bacterial Cells’ (1840301), and ‘Towards a unified theory of regulatory functions and networks across biological and social systems’ (2133863). We thank Jose Miguel Fari ñ a for their helpful comments on the manuscript.

## Notes

### Competing Interest Statement

The authors have declared no competing interest.

## References

1. Marquet, P. A., Navarrete, S. A. & Castilla, J. C. Scaling population density to body size in rocky intertidal communities. Science 250, 1125–1127 (1990).

2. Gillooly, J. F., Brown, J. H., West, G. B., Savage, V. M. & Charnov, E. L. Effects of size and temperature on metabolic rate. science 293, 2248–2251 (2001).

3. West, G. B., Brown, J. H. & Enquist, B. J. A general model for the origin of allometric scaling laws in biology. Science 276, 122–126 (1997).

4. Brown, J. H. & West, G. B. Scaling in biology (Oxford University Press on Demand, 2000).

5. He, J.-H. A brief review on allometric scaling in biology in International Conference on Computational and Information Science (2004), 652–658.

6. Spence, A. J. Scaling in biology. Current Biology 19, R57–R61 (2009).

7. West, G. B. & Brown, J. H. The origin of allometric scaling laws in biology from genomes to ecosystems: towards a quantitative unifying theory of biological structure and organization. Journal of experimental biology 208, 1575–1592 (2005).

8. Ghosh, A. Scaling laws. Mechanics over micro and nano scales, 61–94 (2011).

9. Ribeiro, F. L. & Pereira, W. R. A Gentle Introduction to Scaling Laws in Biological Systems. arXiv preprint arXiv:2105.01540 (2021).

10. Pé rez-Garcia, V. M. et al. Universal scaling laws rule explosive growth in human cancers. Nature physics 16, 1232–1237 (2020).

11. West, G. B., Brown, J. & Enquist, B. Scaling in biology: patterns and processes, causes and consequences. Scaling in biology 87, 112 (2000).

12. Rau, A. Biological scaling and physics. Journal of biosciences 27, 475–478 (2002).

13. Whitfield, J. In the beat of a heart: life, energy, and the unity of nature (Joseph Henry Press, 2006).

14. Antol, A. & Kozlowski, J. Scaling of organ masses in mammals and birds: phylogenetic signal and implications for metabolic rate scaling. ZooKeys 982, 149 (2020).

15. Bettencourt, L. M., Lobo, J., Helbing, D., Kühnert, C. & West, G. B. Growth, innovation, scaling, and the pace of life in cities. Proceedings of the national academy of sciences 104, 7301–7306 (2007).

16. West, G. B., Brown, J. H. & Enquist, B. J. The fourth dimension of life: fractal geometry and allometric scaling of organisms. science 284, 1677–1679 (1999).

17. Brown, J. H., Gillooly, J. F., Allen, A. P., Savage, V. M. & West, G. B. Toward a metabolic theory of ecology. Ecology 85, 1771–1789 (2004).

18. Carscadden, K. A. et al. Niche breadth: causes and consequences for ecology, evolution, and conservation. The Quarterly Review of Biology 95, 179–214 (2020).

19. Ducatez, S., Sol, D., Sayol, F. & Lefebvre, L. Behavioural plasticity is associated with reduced extinction risk in birds. Nature Ecology & Evolution 4, 788–793 (2020).

20. Roughgarden, J. Evolution of niche width. The American Naturalist 106, 683–718 (1972).

21. Futuyma, D. J. & Moreno, G. The evolution of ecological specialization. Annual review of Ecology and Systematics 19, 207–233 (1988).

22. Poisot, T., Bever, J. D., Nemri, A., Thrall, P. H. & Hochberg, M. E. A conceptual framework for the evolution of ecological specialisation. Ecology letters 14, 841–851 (2011).

23. Brändle, M., Prinzing, A., Pfeifer, R. & Brandl, R. Dietary niche breadth for Central European birds: correlations with species-specific traits. Evolutionary Ecology Research 4, 643–657 (2002).

24. Gaston, K. J. & Spicer, J. I. The relationship between range size and niche breadth: a test using five species of Gammarus (Amphipoda). Global Ecology and Biogeography 10, 179–188 (2001).

25. Slatyer, R. A., Hirst, M. & Sexton, J. P. Niche breadth predicts geographical range size: a general ecological pattern. Ecology letters 16, 1104–1114 (2013).

26. Hackett, S. J. et al. A phylogenomic study of birds reveals their evolutionary history. science 320, 1763–1768 (2008).

27. Oro, D., Cam, E., Pradel, R. & Martınez-Abraın, A. Influence of food availability on demography and local population dynamics in a long-lived seabird. Proceedings of the Royal Society of London. Series B: Biological Sciences 271, 387–396 (2004).

28. Shealer, D. A. Foraging behavior and food of seabirds. Biology of marine birds 14, 137–177 (2002).

29. De Graaf, R. M., Tilghman, N. G. & Anderson, S. H. Foraging guilds of North American birds. Environmental Management 9, 493–536 (1985).

30. Gonzá lez-Salazar, C., Martınez-Meyer, E. & Ló pez-Santiago, G. A hierarchical classification of trophic guilds for North American birds and mammals. Revista Mexicana de Biodiversidad 85, 931–941 (2014).

31. Costa, G. C. Predator size, prey size, and dietary niche breadth relationships in marine predators. Ecology 90, 2014–2019 (2009).

32. Davis, R. B., Õ unap, E., Javoi š, J., Gerhold, P. & Tammaru, T. Degree of specialization is related to body size in herbivorous insects: a phylogenetic confirmation. Evolution 67, 583–589 (2013).

33. Ballance, L., Ainley, D. G. & Hunt Jr, G. L. Seabird foraging ecology. Encyclopedia of ocean sciences 5, 2636–2644 (2001).

34. Wawrzynek-Borejko, J., Panasiuk, A., Hinke, J. T. & Korczak-Abshire, M. Are the diets of sympatric Pygoscelid penguins more similar than previously thought? Polar Biology 45, 1559–1569 (2022).

35. Bradstreet, M. S. & Brown, R. G. Feeding ecology of the Atlantic Alcidae. (1985).

36. Hedd, A. & Montevecchi, W. A. Diet and trophic position of Leach’s storm-petrel Oceanodroma leucorhoa during breeding and moult, inferred from stable isotope analysis of feathers. Marine Ecology Progress Series 322, 291–301 (2006).

37. Barrett, R. T. et al. Diet studies of seabirds: a review and recommendations. ICES Journal of Marine Science 64, 1675–1691 (2007).

38. Voje, K. L., Hansen, T. F., Egset, C. K., Bolstad, G. H. & Pé labon, C. Allometric constraints and the evolution of allometry. Evolution 68, 866–885 (2014).

39. Pé labon, C. et al. Evolution of morphological allometry. Annals of the New York Academy of Sciences 1320, 58–75 (2014).

40. Kim, S. et al. Comparison of carnivore, omnivore, and herbivore mammalian genomes with a new leopard assembly. Genome biology 17, 1–12 (2016).

41. Gillooly, J. F., McCoy, M. W. & Allen, A. P. Effects of metabolic rate on protein evolution. Biology letters 3, 655–660 (2007).

42. Omland, K. E. Correlated rates of molecular and morphological evolution. Evolution 51, 1381–1393 (1997).

43. Seligmann, H. Positive correlations between molecular and morphological rates of evolution. Journal of Theoretical Biology 264, 799–807 (2010).

44. Creighton, M. J., Greenberg, D. A., Reader, S. M. & Mooers, A. Ø. The role of behavioural flexibility in primate diversification. Animal Behaviour 180, 269–290 (2021).

45. Macdonald, D. The encyclopedia of mammals (OUP Oxford, 2009).

