## Supplementary Table 1 for "Foraging specialization and body size in seabirds"

Material

| Table S1. Parameters for alternative analysis | | | | |
|---|---|---|---|---|
| Relationship | Parameters | | | |
| Not averaging | slope | intercept | r-sq. | p |
| Diets-size | -0.00695 | 0.5453 | 0.0003 | 0.73 |
| Strategies-size | -0.1699 | 0.8143 | 0.1394 | 2.689*10^11 |
| Strategies-diet | 0.308 | | 0.14 | 2.02*10^-12 |
| Diets/strategy-size | -0.1591 | | 0.1152 | 2.31*10^-9 |
| Averaging diet/strategies | | | | |
| ‹Diets›-size interval (100 g) | -0.14±0.05 | 1.02 | 0.12 | 0.006 |
| ‹Strategies›-size interval (100 g) | -0.19±0.06 | 0.93 | 0.15 | 0.002 |
| Strategies-diet (for intervals body size of 100 g) | 0.0707 | 1.8123 | 0.0006 | 0.55 |
| ‹Diets/Strategies›-size interval (100 g) | -0.1114 | | 0.035 | 0.15 |
